# Spelling changes and fluorescent tagging with prime editing vectors for plants

**DOI:** 10.1101/2020.07.16.206276

**Authors:** Li Wang, Hilal Betul Kaya, Ning Zhang, Rhitu Rai, Matthew R. Willmann, Sara C. D. Carpenter, Andrew C. Read, Federico Martin, Zhangjun Fei, Jan E. Leach, Gregory B. Martin, Adam J. Bogdanove

## Abstract

Prime editing (PE) is a recent adaptation of the CRISPR-Cas system that uses a Cas9(H840A)-reverse transcriptase (RT) fusion and a guide RNA (pegRNA) amended with template and primer binding site (PBS) sequences to achieve RNA-templated conversion of the target DNA, allowing specified substitutions, insertions, and deletions. In the first report of PE in plants, a variety of edits in rice and wheat were described, including insertions up to 15 bp. Several studies in rice quickly followed, but none reported a larger insertion. Here, we report easy-to-use vectors for PE in dicots and monocots, their validation in *Nicotiana benthamiana*, rice and Arabidopsis, and an insertion of 66 bp that enabled split-GFP fluorescent tagging.

---

Prime editing (PE) is a recent adaptation of the CRISPR-Cas system that uses a Cas9(H840A)-reverse transcriptase (RT) fusion and a guide RNA (pegRNA) amended with template and primer binding site (PBS) sequences to achieve RNA-templated conversion of the target DNA, allowing specified substitutions, insertions, and deletions (Anzalone et al., 2019). The largest insertion reported to date was in human cells, a 44-bp *loxP* tag (Anzalone et al., 2019). In the first report of PE in plants, a variety of edits in rice and wheat were described, including insertions up to 15 bp (Lin et al. 2020). Several studies in rice quickly followed, but none reported a larger insertion. Here, we report easy-to-use vectors for PE in dicots and monocots (**Figure 1a**), their validation in three plant species, and an insertion of 66 bp that enabled split-GFP fluorescent tagging.

**Figure 1.**
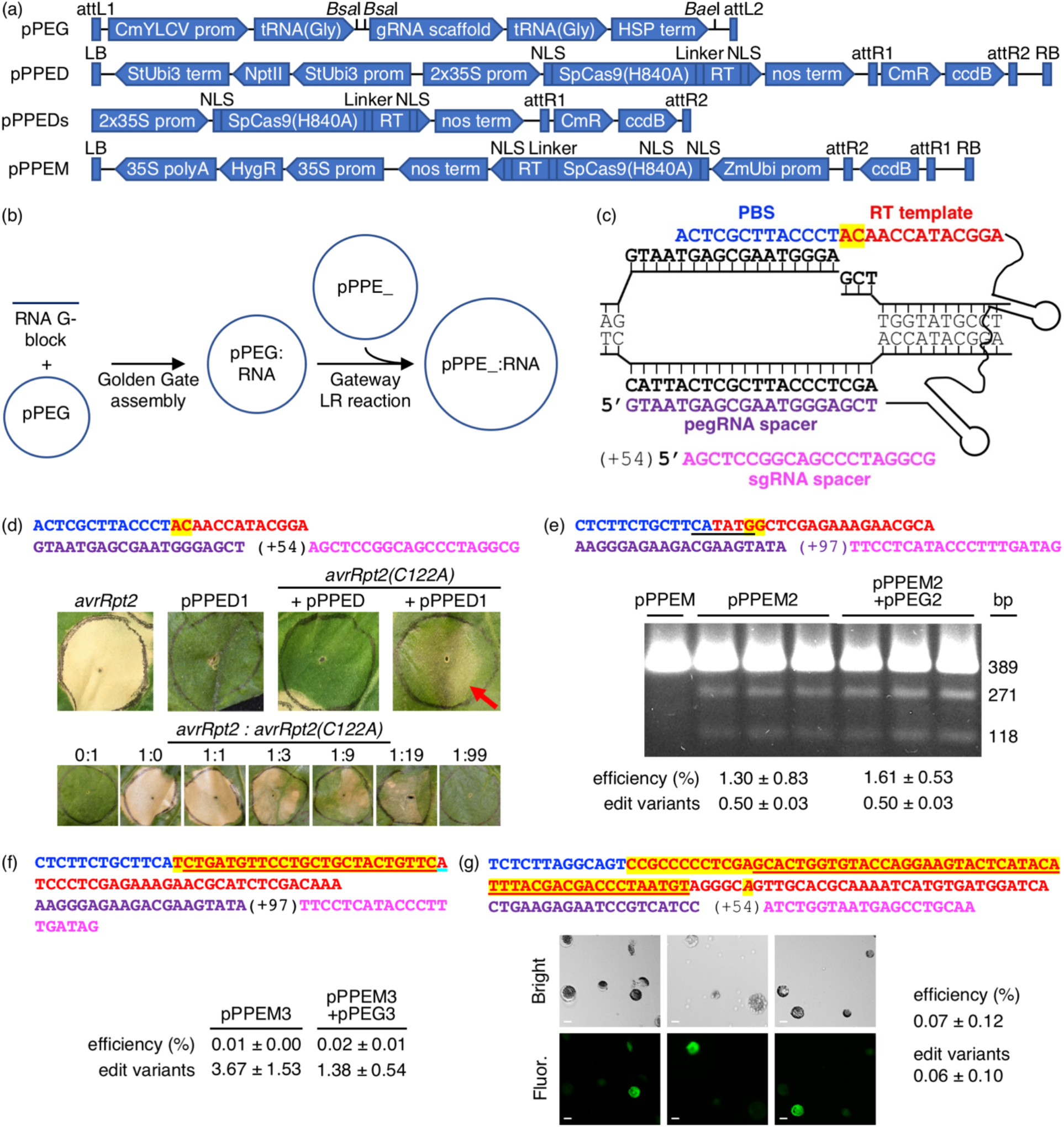
Vectors for PE in plants and their use for spelling changes and tag insertions. **(a)** The expression and cloning cassettes in each vector (vectors and details available at www.addgene.org). **(b)** Workflow for generating PE constructs. **(c)** A pegRNA on its target, showing the PBS (blue), RT template (red, with the edit highlighted in yellow), spacer (purple), and pegRNA-directed nick, and, for PE3, an sgRNA (magenta) with the position of its nick site relative to the pegRNA nick in parentheses. **(d)** A 2-bp edit of *avrRpt2(C122A)* using agroinfiltration of *Nicotiana benthamiana* leaves. Top, features of the pegRNA and sgRNA, colored as in (c). Middle, leaves 12 days after introduction of *avrRpt2* (OD_600_=0.05), or pPPED1, *avrRpt2(C122A)* and empty pPPED, or *avrRpt2(C122A)* and pPPED1 (each at final OD_600_=0.5). Bottom, leaves 6 days after infiltration of different ratios of strains delivering *avrRpt2* or *avrRpt2(C122A)*, both at OD_600_=0.05 before mixing. **(e)** A 2-bp substitution at *OsSULTR3;6* in rice protoplasts. Top, pegRNA and sgRNA features with *Bst*Z17I site underlined. Below, *Bst*Z17I digested amplicons from protoplasts transfected with empty pPPEM, pPPEM2, or pPPEM2 and equimolar pPEG2. **(f)** A 25-bp insertion for FLAG tagging at *OsSULTR3;6*. Top, pegRNA and sgRNA features, with FLAG coding sequence underlined. **(g)** A 60-bp insertion for fusion of GFP11 at AT1G26660.1 in Arabidopsis CYTO-sfGFP1-10^OPT^ protoplasts. Top, pegRNA and sgRNA features, with GFP11 coding sequence underlined and a substitution in the pegRNA PAM italicized. Below, bright field and fluorescence micrographs of protoplasts from replicate transfections; scale bar, 10 μm. Bottom of **e-g**, editing efficiencies determined by amplicon deep-sequencing adjusted for transfection efficiency, and, relative to the number of perfect edit reads set as 1, the number of variant reads with the edit but not included in calculated efficiency (see text). Amplicon sequences were analyzed using CRISPResso2 (Clement et al., 2019) and are available under NCBI BioProject PRJNA641949.

We designed the vectors for PE2, which incorporates an engineered RT, or PE3, which adds an sgRNA to nick the non-edited strand and drive its conversion (Anzalone et al., 2019). For dicots we replaced *Cas9* in binary vector p201N (Cermak et al., 2017) with *Cas9(H840A)* from pMOD_A0301 (Cermak et al., 2017) plus a tomato codon-optimized linker and engineered RT sequence (Anzalone et al., 2019), then added a Gateway destination cassette (Thermo-Fisher), creating pPPED. Also, for transfection or bombardment, we created pPPEDs, using pBluescript KS(-). For monocots, we mutated pUbi-Cas9 (Zhou et al., 2014) to encode Cas9(H840A), then added the linker and RT sequence, optimized for rice, yielding pPPEM. We created an entry vector for the RNA modules, pPEG, by inserting a CmYLCV promoter-driven cassette with *BsaI* cloning sites followed by a gRNA scaffold, together flanked by tRNA (Gly) sequences, and a unique *BaeI* site downstream for additional elements, into pCR8/GW/TOPO (Thermo-Fisher). To prepare a construct, pegRNA sequence without scaffold (PE2), or pegRNA with scaffold followed by tRNA(Gly) and an sgRNA spacer (PE3), with a *Bsa*I site and compatible sequence at each end, is synthesized and introduced by Golden Gate reaction (Engler et al., 2008) into pPEG; the resulting module is transferred by LR recombination into pPPED, pPPEDs, or pPPEM (**Figure 1b**).

Our editing strategy was PE3. Example peg- and sgRNAs are shown in **Figure 1c**. First, we tested pPPED by agroinfiltration of *Nicotiana benthamiana* leaves (**Figure 1d**). The edit was a GC to TG “correction” of a mutation in the *avrRpt2* gene of *Pseudomonas syringae (avrRpt2[C122A];* Mazo-Molina et al., 2020), delivered by a co-infiltrated strain, that would restore the gene’s ability to elicit plant cell death. Together with *avrRpt2(C122A)*, pPPED carrying a pegRNA/sgRNA module for the edit (pPPED1), but not empty pPPED and not pPPED1 alone, resulted in cell death. To estimate efficiency, we determined the sensitivity of the assay by co-infiltrating different ratios of *avrRpt2* and *avrRpt2(C122A)* strains. The *avrRpt2* strain was sufficient for cell death at OD_600_=0.0025 (1:19) but not at OD_600_=0.0005 (1:99). Thus, in the editing experiment, in which the *avrRpt2(C122A)* strain was at OD_600_=0.5, more than 0.1% (0.0005/0.5) and likely 0.5% (0.0025/0.5) or more of the delivered *avrRpt2(C122A)* was converted to wild type. Amplicon deep sequencing detected only 0.06% (± 0.03%, four infiltrations), likely because the template included *avrRpt2(C122A)* on the vector in *Agrobacterium*, not exposed to the PE reagent.

Next, we tested pPPEM in rice (cv. Nipponbare) protoplasts, on a chromosomal target, *OsSULTR3;6* (LOC_Os01g52130; Cernadas et al., 2014) (**Figure 1e**). The edit, GG to CC, eliminates the stop codon and introduces a *Bst*Z17I site. In three transfections with pPPEM carrying the pegRNA/sgRNA module for the edit (pPPEM2), but not in a control transfection with empty pPPEM, *Bst*Z17I digestion of PCR product spanning the target confirmed editing. Amplicon sequencing revealed efficiencies ranging from 0.7% to 2.2%, when adjusted for transfection efficiency (~41%). An equimolar amount of entry vector carrying the RNA module, pPEG2, added to the pPPEM2 transfections did not increase average editing efficiency (unpaired, one tail *t*-test, p<0.05).

Third, again using pPPEM in rice protoplasts and targeting the same location in *OsSULTR3;6*, we attempted a 25-bp insertion for translational fusion of the FLAG epitope (**Figure 1f**). We carried out three transfections with the editing construct, pPPEM3, and three more with the corresponding pPEG plasmid, pPEG3, added. Amplicon sequencing confirmed insertion, but at relatively low adjusted efficiency, not significantly altered by pPEG3 (0.02 ± 0.01% and 0.01 ± 0.00%, respectively).

Finally, we tested pPPEDs in protoplasts of Arabidopsis lines expressing ß strands 1-10 of optimized super-fold green fluorescent protein targeted to the cytoplasm, CYTO-sfGFP1-10^OPT^, or nucleus, NUC-sfGFP1-10^OPT^ (Park et al., 2017). The edit was a 66-bp insertion encoding a linker and GFP11. We reasoned that a split-GFP approach could enable fluorescent tagging despite the apparent insertion size limitation of PE. Indeed, CYTO-sfGFP1-10^OPT^ transfections with a pPPEDs construct, pPPEDs4, targeting the insertion to the cytosolic prefoldin chaperone subunit family protein gene AT1G26660.1 yielded fluorescent protoplasts (**Figure 1g**), while control transfections of NUC-sfGFP1-10^OPT^ with the same construct, or of CYTO-sfGFP1-10^OPT^ with a pPPEDs construct, pPPEDs5, targeting the insertion to the histone 2B gene (AT5G22880), did not. The pPPEDs4 RT template includes a C to A substitution 6 bp after the GFP11 sequence that destroys the pegRNA PAM, a strategy proposed to limit indel formation between the PE3 nicks and to disfavor reversion of the edited strand (Anzalone et al., 2019). Amplicon sequencing of the CYTO- and NUC-sfGFP1-10^OPT^ transfections with pPPEDs4 (three each) confirmed successful insertion, averaging 0.07±0.12% adjusted efficiency.

For all edits, the positive amplicon reads included some with other differences from wild type in the window encompassing the nick sites and edit plus 2 bp on either side, and some of the insertion edit reads had one or more SNPs or indels in the insertion. These types of reads were not counted in the reported efficiencies. They may represent unintended PE, spontaneous mutations, or artifact. Sequencing of the AT1G26660.1 amplicon from the pPPEDs5 negative control transfection, and from a transfection of NUC-GFP1-10^OPT^ with pPPEDs5, yielded an average of 6.4±4.0% reads varying from wild type, so editing efficiencies may have been higher than we calculated counting only perfect reads.

In summary, we developed vectors for straightforward plant PE construct assembly and demonstrated their efficacy in one monocot and two dicot species. Edits included two 2-bp codon changes, a 24-bp FLAG tag insertion, and a 66-bp GFP11 insertion. The 66-bp insertion is the largest reported for PE and provides important proof of concept for fluorescent tagging using PE. Editing efficiencies, especially for insertions, were low. However, efficiencies are likely to be higher in stably transformed plants or with meristem transformation (Maher et al., 2020), and the vectors thus useful in extending PE to diverse plant species.

## Acknowledgements

This work was supported by the National Science Foundation (IOS-1444511 to AB and JL, and IOS-1546625 to GM), the National Institute of Food and Agriculture of the U.S. Department of Agriculture (2018-67011-28025 to AR), the Indo-U.S. Science and Technology Forum, Government of India (fellowship to RR), and Manisa Celal Bayar University Scientific Research Projects funds (2017-113 to HK). Fluorescence microscopy was performed at the Imaging Facility of the Biotechnology Resource Center at Cornell University’s Institute of Biotechnology, which was supported by the National Institutes of Health (S10RR025502).

## References

Anzalone, A.V., Randolph, P.B., Davis, J.R., Sousa, A.A., Koblan, L.W., Levy, J.M., Chen, P.J., Wilson, C., Newby, G.A., Raguram, A. and Liu, D.R. (2019) Search-and-replace genome editing without double-strand breaks or donor DNA. Nature 576, 149–157.

Cermak, T., Curtin, S.J., Gil-Humanes, J., Cegan, R., Kono, T.J.Y., Konecna, E., Belanto, J.J., Starker, C.G., Mathre, J.W., Greenstein, R.L. and Voytas, D.F. (2017) A multipurpose toolkit to enable advanced genome engineering in plants. Plant Cell 29, 1196–1217.

Cernadas, R.A., Doyle, E.L., Nino-Liu, D.O., Wilkins, K.E., Bancroft, T., Wang, L., Schmidt, C. L., Caldo, R., Yang, B., White, F.F., Nettleton, D., Wise, R.P. and Bogdanove, A.J. (2014) Code-assisted discovery of TAL effector targets in bacterial leaf streak of rice reveals contrast with bacterial blight and a novel susceptibility gene. PLoS Path. 10, e1003972.

Clement, K., Rees, H., Canver, M.C., Gehrke, J.M., Farouni, R., Hsu, J.Y., Cole, M.A., Liu, D. R., Joung, J.K., Bauer, D.E. and Pinello, L. (2019) CRISPResso2 provides accurate and rapid genome editing sequence analysis. Nat. Biotechnol. 37, 224–226.

Engler, C., Kandzia, R. and Marillonnet, S. (2008) A one pot, one step, precision cloning method with high throughput capability. PLoS ONE 3, e3647.

Maher, M.F., Nasti, R.A., Vollbrecht, M., Starker, C.G., Clark, M.D. and Voytas, D.F. (2020) Plant gene editing through *de novo* induction of meristems. Nat. Biotechnol. 38, 84–89.

Mazo-Molina, C., Mainiero, S., Haefner, B.J., Bednarek, R., Zhang, J., Feder, A., Shi, K., Strickler, S.R. and Martin, G.B. (2020) *Ptr1* evolved convergently with *RPS2* and *Mr5* to mediate recognition of AvrRpt2 in diverse solanaceous species. Plant J.

Park, E., Lee, H.Y., Woo, J., Choi, D. and Dinesh-Kumar, S.P. (2017) Spatiotemporal monitoring of *Pseudomonas syringae* effectors via type III secretion using split fluorescent protein fragments. Plant Cell 29, 1571–1584.

Zhou, H., Liu, B., Weeks, D.P., Spalding, M.H. and Yang, B. (2014) Large chromosomal deletions and heritable small genetic changes induced by CRISPR/Cas9 in rice. Nucleic Acids Res. 42, 10903–10914.

